# Genetic diversity and mother-child overlap of the gut associated microbiota determined by reduced genome sequencing

**DOI:** 10.1101/191445

**Authors:** Anuradha Ravi, Ekaterina Avershina, Inga Leena Angell, Jane Ludvigsen, Prashanth Manohar, Sumathi Padmanaban, Ramesh Nachimuthu, Knut Rudi

**Affiliations:** Faculty of Chemistry, Biotechnology and Food Science, Norwegian University of Life Sciences, Ås, Norway; Antibiotic resistance and phage therapy laboratory, Department of Biomedical sciences, School of Bio-sciences and technology, VIT University, Tamil Nadu, India; Nishanth Hospital, Tamil Nadu, India

## Abstract

The genetic diversity and sharing of the mother-child associated microbiota remain largely unexplored. This severely limits our functional understanding of gut microbiota transmission patterns. The aim of our work was therefore to use a novel reduced metagenome sequencing in combination with shotgun and 16S rRNA gene sequencing to determine both the metagenome genetic diversity and the mother-to-child sharing of the microbiota. For a cohort of 17 mother-child pairs we found an increase of the collective metagenome size from about 100 Mbp for 4-day-old children to about 500 Mbp for mothers. The 4-day-old children shared 7% of the metagenome sequences with the mothers, while the metagenome sequence sharing was more than 30% among the mothers. We found 15 genomes shared across more than 50% of the mothers, of which 10 belonged to *Clostridia*. Only *Bacteroides* showed a direct mother-child association, with *B. vulgatus* being abundant in both 4-day-old children and mothers. In conclusion, our results support a common pool of gut bacteria that are transmitted from adults to infants, with most of the bacteria being transmitted at a stage after delivery.

## INTRODUCTION

The colonization by gut bacteria at infancy is crucial for proper immune development and gut maturation (1). At birth, we are nearly sterile, while just after a few days of life we become densely colonized by bacteria (2). How and when we acquire the adult associated bacteria,are not yet completely established (2). Recent 16S rRNA gene sequence data suggest that most of the adult associated bacteria are recruited at a stage after delivery (3), while shotgun analyses suggest a high frequency of direct transmission during delivery (4). The limitations of these studies, however, are that 16S rRNA gene analyses do not have sufficient resolution to resolve mother to child transmission at the strain level, while shotgun sequencing requires extensive and complex analyses (4,5). Taken together, this restricts the possibility of gaining broad-scale knowledge about the microbiota genetic diversity and distribution with the current analytical approaches. There is thus a need for analytical approaches that combine efficiency and resolution.

The aim of the current work was therefore to use a novel concept of reduced metagenome sequencing (RMS; schematically outlined in Fig. 1) in combination with 16S rRNA gene and shotgun sequencing to estimate genetic diversity and mother-child overlap for gut associated bacteria for a medium size cohort of 17 mother-child pairs.

**Figure 1.**
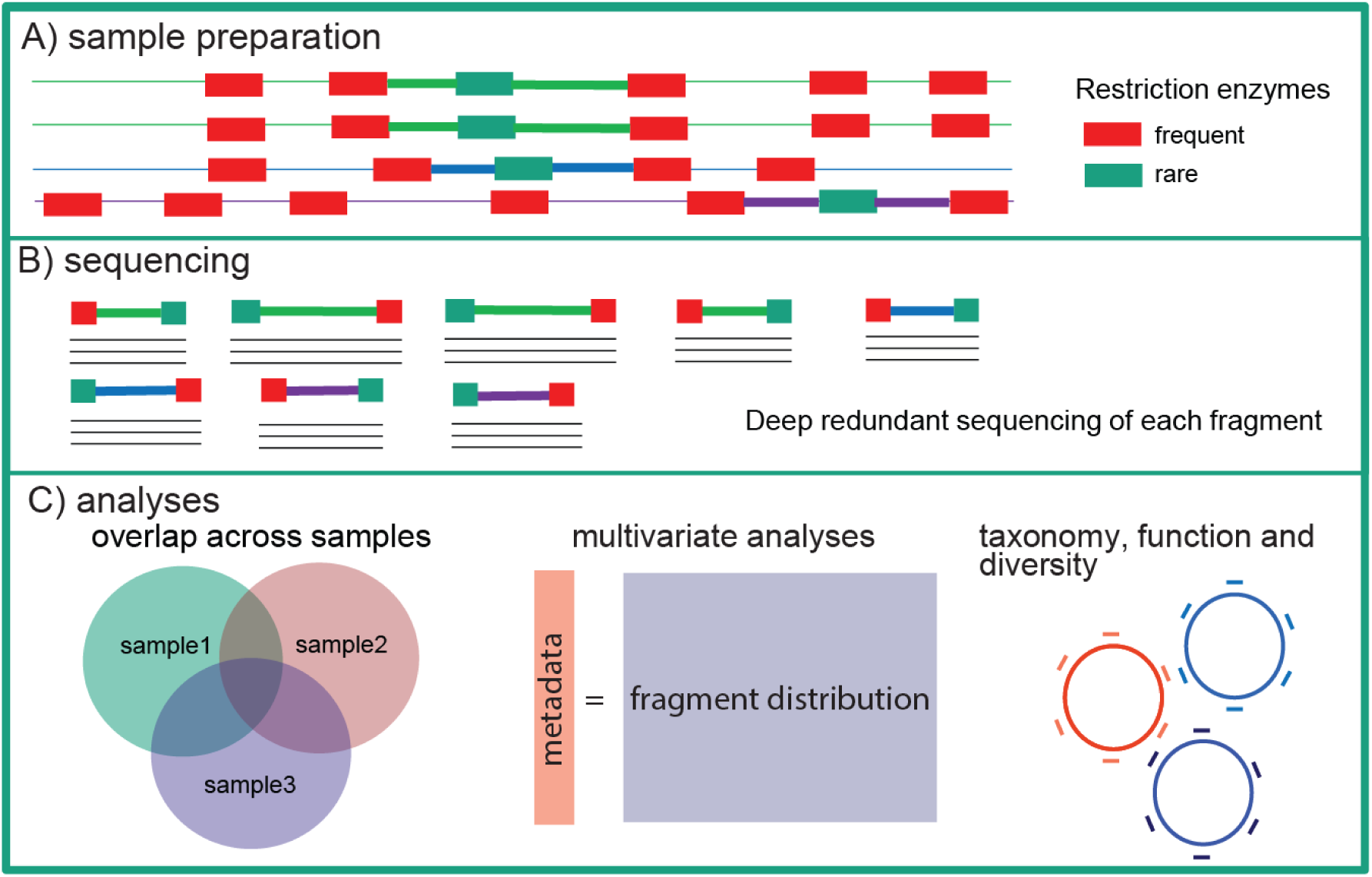
Schematic outline of the reduced metagenome sequencing approach. (A) In the first stage we amplify the fragments flanked by a frequent and a rare restriction enzyme cutting sites by the RMS principle (indicated by thick lines). (B) The amplified fragments are then sequenced by deep redundant sequencing with high coverage for each fragment. (C) The fragment frequency information can be used for different analytical applications. The overlap in fragments across samples can be used as a proxy for high resolution analyses of the overlap in microbiota. Fragment frequencies can be directly related to metadata. Finally, fragment distribution can be used to estimate taxonomy, function and genetic diversity of the microbiota.

RMS requires a relatively shallow sequencing depth in order to gain insight into genetic diversity and microbiota overlap across individuals. With this approach only a defined fraction of the metagenome is sequenced. Principles related to reduced metagenome sequencing have been widely used for strain resolution analyses since the 1990s by DNA fragment size separation (6,7). RMS, however, gives additional information about fragment sequence, therefore thus enabling the estimation of total genetic diversity and overlap of metagenome sequences. Thus, RMS has the potential to solve some of the most urgent needs in current metagenome sequence analyses.

Our study presents evidence that less than 10% of the microbiota is directly transmitted from mother to child, while the sharing of more than 30% of the microbiota across random mothers suggests that the majority of the gut microbiota is transmitted at a stage after delivery.

## MATERIALS AND METHODS

### Cohort description

The study consists of an unselected longitudinal cohort of 17 mother-infant pairs. The infants were born full term at Nishanth Hospital, India. Twelve of the 17 infants were born through cesarean section. All the mothers that gave birth through vaginal or cesarean section were given antibiotics either during pregnancy, during labor and/or after pregnancy (Suppl. Table 1). Fecal samples were collected from late pregnant women (gestational age 32-36 weeks) and in infants 0-4 days after birth for all the mother child-pairs, while the numbers of samples at 15 days were 4, 60 days 3 and 120 days 3. Fecal samples were collected and stored at -20°C up to a week with STAR buffer (Roche, Basel, Switzerland). Then, the samples were transferred to -80°C for longer storage. One of the parents of each child signed a written informed consent form before the fecal sample collection, which is in accordance with legislation in India.

### Mock community

Mock communities of *E.coli* ATCC25922; *E. faecalis* V583; *B. longum* DSM20219 and *B. infantis* DSM20088 mixed in varying proportions ranging from 0% to 100% were used to assess sensitivity of the AFLP sequencing in prediction and identification of bacterial strains.

### DNA isolation

The fecal samples were diluted 3-fold with STAR-buffer and pre-centrifuged at 1200 rpm for 8 sec to remove large particles. An overnight culture of each of the four bacteria species used for mock community analyses was used for DNA isolation. Bacterial cultures were pre-centrifuged at 13000 rpm for 5 min, and then pellets were washed twice in 1x PBS buffer. The supernatant from the pre-centrifuged stool samples, as well as bacterial pellets in PBS, were mixed with acid-washed glass beads (Sigma-Aldrich, <106μm; 0.25g) and bead-beated at 1800rpm in 40 seconds twice, with 5 minutes’ rest between the runs. The samples were then centrifuged at 13000rpm for 5 minutes. An automated protocol based on paramagnetic particles (LGC Genomics, UK) was used for the DNA isolation using a KingFisher Flex (ThermoFisher Scientific, USA), following the manufacturers recommendations. After extraction, samples were quantified and normalized using Qubit fluorometer (ThermoFisher Scientific, USA). The DNA concentrations were normalized to 0.2 ng/μl prior to further processing. Mock communities were prepared to contain a total of 10 ng DNA in each sample.

### Library preparation and sequencing

For RMS, DNA fragments were obtained by cutting genomic DNA using an enzyme combination of EcoRI and MseI, followed by an adapter ligation and PCR amplification. Restriction cutting was performed in 20 μl volumes containing 8U EcoRI (New England Biolabs, USA), 4U MseI (New England Biolabs, USA), 1x Cut smart buffer (New England Biolabs, USA), and 1 ng genomic DNA. The samples were incubated at 37°C for one hour to make sure that the restriction enzymes would cut appropriate amounts of DNA into fragments. For PCR amplification of the fragments, adapters were ligated onto the fragments. This was done by adding the sample a 5 μl volume of 0.5 μM EcoRI adapter mix, 5μM MseI adapter mix, 1 μl T4 DNA ligase (New England Biolabs, USA) and 1x T4 reaction buffer (New England Biolabs, USA). The adapter mixes were made of equal volumes of forward (EcoRI; 5’- CTCGTAGACTGCGTACC-3’, MseI; 5’- GACGATGAGTCCTGAG-3’) and reverse adapters (EcoRI; 5’- AATTGGTACGCAGTCTAC-3’, MseI; 5’- TACTCAGGACTCAT-3’). The adapters will ligate to the fragments that have been cut by both restriction enzymes. The samples were incubated for 3 hours at 37°C. PCR amplification was performed using primer pairs EcoRI (5’- GACTGCGTACCAATTC-3’)/MseI (5’- GATGAGTCCTGAGTAA-3’), targeting the RMS fragments, and PRK341F (5’- CCTACGGGRBGCASCAG-3’)/ PRK806R(5’- GGACTACYVGGGTATCTAAT-3’) (8),targeting the V3-V4 region of the 16 S rRNA gene. Each reaction contained 1x HotFirePol DNA polymerase Ready to load (Solis BioDyne, Estonia), 0.2μM forward and reverse primer (Invitrogen, USA) and 2 μl template DNA. The cycling conditions for the 16S rRNA gene were 25 cycles of 95°C for 30 seconds, 55°C for 30 seconds and 72°C for 45 seconds, while the cycling conditions for the RMS were 25 cycles of 95°C for 30 sec, 56°C for 1 min and 72°C for 1 min. The PCR products were purified by AMPure XP beads (Beckman Coulter, USA) prior to further processing.

To index the fragments 1x FirePol DNA polymerase Ready to load (Solis BioDyne, Estonia), 0.2μM forward index primer and reverse index primer and 2μl purified PCR products were used. The fragments were amplified by PCR using the following thermal cycle: 95° C for 5 minutes, followed by 10 cycles of 95° C for 30 seconds, 55° C for 60 seconds and 72° C for 45 seconds. A final step at 72° C for 7 minutes was included. All samples were pooled using 100ng DNA from each sample. The pooled sample were then purified using AMPure XP beads prior to sequencing. The pooled sequencing library comprised 15% PhiX and 85% pooled sample.

The shotgun metagenome sequencing was done as previously described using Illumina Nextera XT, following the recommendations by the producer (9).

### Data analysis

Raw data of 16S were analyzed using a standard workflow of QIIME pipeline (3). Sequences were paired-end joined (join_paired_ends.py with *fastq_join* method), demultiplexed using split_library.py script with no error in the barcode allowed (max_barcode_errors 0) and barcodes removed. Sequences were then filtered using *fastq_filter* command of *usearch* (maxee = 0.22; minlen = 350). Finally, sequences were clustered at 97 % similarity threshold using *cluster_otus* command of *usearch*. Singletons were removed and an additional reference-based chimera removal step against GOLD database was performed. Resulting dataset was then rarefied to 6000 sequences per sample (Schematically outlined in Suppl. Fig 1A). The reduced metagenome and shotgun data were analyzed using a CLC Genomic Workbench (Qiagen, Hilden, Germany), using the paired-end sequence-merging tool, *de novo* and reference based assembly tools, and Blast searches (Schematically outlined in Suppl. Fig. 1B). The shotgun data were not processed further after *de novo* assembly, while for the RMS analyses we followed the bioinformatics workflow, as outlined in Fig. 2. Sample comparisons and statistical analyses were done using Matlab 2016a (Mathworks Inc, USA) with the PLS toolbox module (Eigenvector Research Inc., USA).

**Figure 2.**
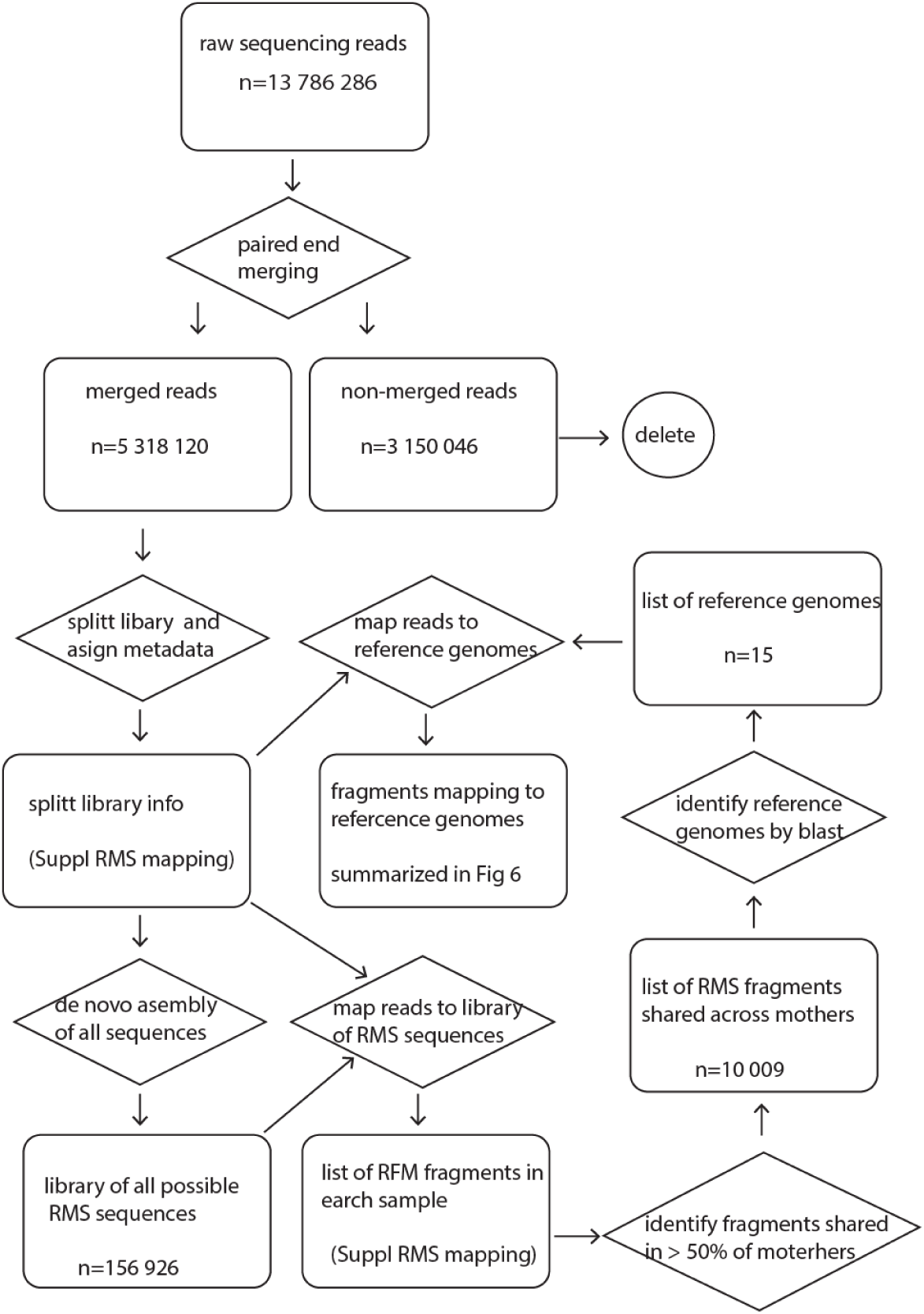
Bioinformatics workflow for the RMS analyses. The workflow is summarized by outputs in rounded squares, while the processes are illustrated with polygons. Directions of processes are illustrated by arrows.

## RESULTS

### Metagenome sequence size

We identified 156 926 unique RMS fragments for the samples analyzed (Fig. 2). This corresponds to complete metagenome sequence size of approximately 750 Mbp given RMS fragment sizes of each 5000 bp. We found an approximately 5-fold increase in the collective metagenome sequence size when comparing 4-day-old children with the mothers (Fig. 3A). The collective metagenome sequence for the 4-day-old children was estimated to approximately 100 Mbp with 40±6.5 Mbp per individual, while the estimated collective metagenome sequence size for mothers was 500 Mbp – with individual sizes of 100 ±7 Mbp.

**Figure 3.**
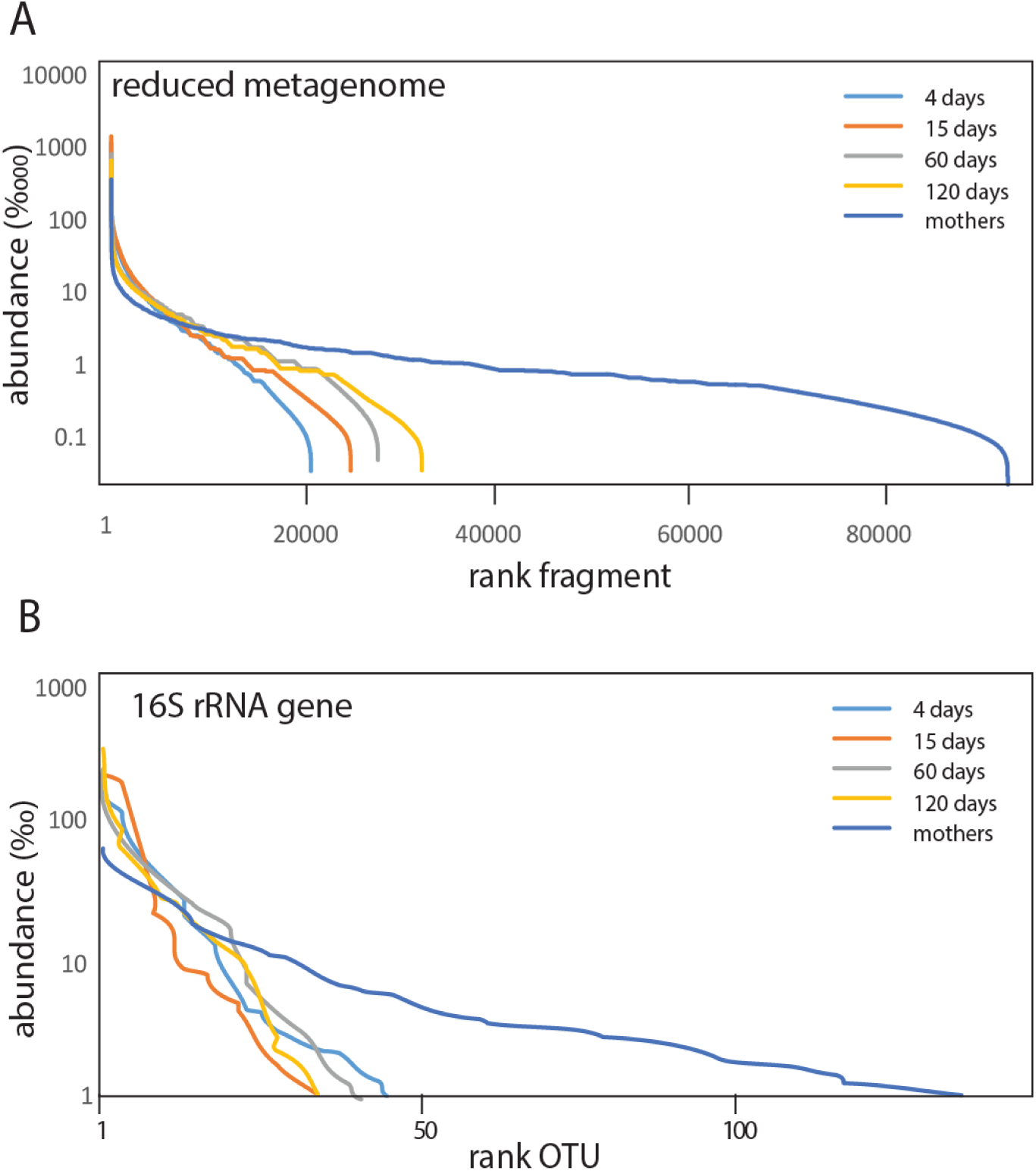
Rank relative abundance distribution across age for (A) RMS fragments and (B) 16S rRNA gene OTUs.

The total number of 16S rRNA gene derived OTUs was 458. For the 4-day-old children we detected a total of 40 OTUs with mean relative abundance > 1 ‰ (13.1±4 per individual), while the number increased to 140 OTUs for the mothers (70.2±7.5 per individual) (Fig. 3B).

For the shotgun analyses we generated 8.6 million paired end reads with a total size 2 070 Mbp shotgun metagenome sequence data for the 4-day-old children, and 6.7 million reads with a total of 1 3964 Mbp sequence for the mothers. However, the shotgun sequences only generated 15.8 Mbp assembly for the 4-day-old children, and 4.3 Mbp for the mothers (Suppl. Table 2).

### Associations of microbiome with mode of delivery and antibiotic usage

For the 4-day-old children we found 27 reduced metagenome sequencing fragments unique to children delivered by c-section, while 20 fragments were unique to the children delivered vaginally (Suppl. Table 3). The most pronounced differences were an overrepresentation of fragments related to the genus *Bacteroides* for vaginal delivery (p=0.0038, Binominal test).

Based on ResFinder assignments (10) of the RNS fragments, we identified 13 fragments associated with known antibiotic resistance genes (Suppl. Table 4). There was a clear association between antibiotic usage during labor and antibiotic resistance genes (p=0.001, ASCA-ANOVA), with Fosfomycin, Beta-lactam and Phenicol resistance showing positive associations (Suppl. Fig 4). Antibiotic usage during pregnancy or after delivery, however, did not seem to affect resistance gene composition (results not shown).

**Figure 4.**
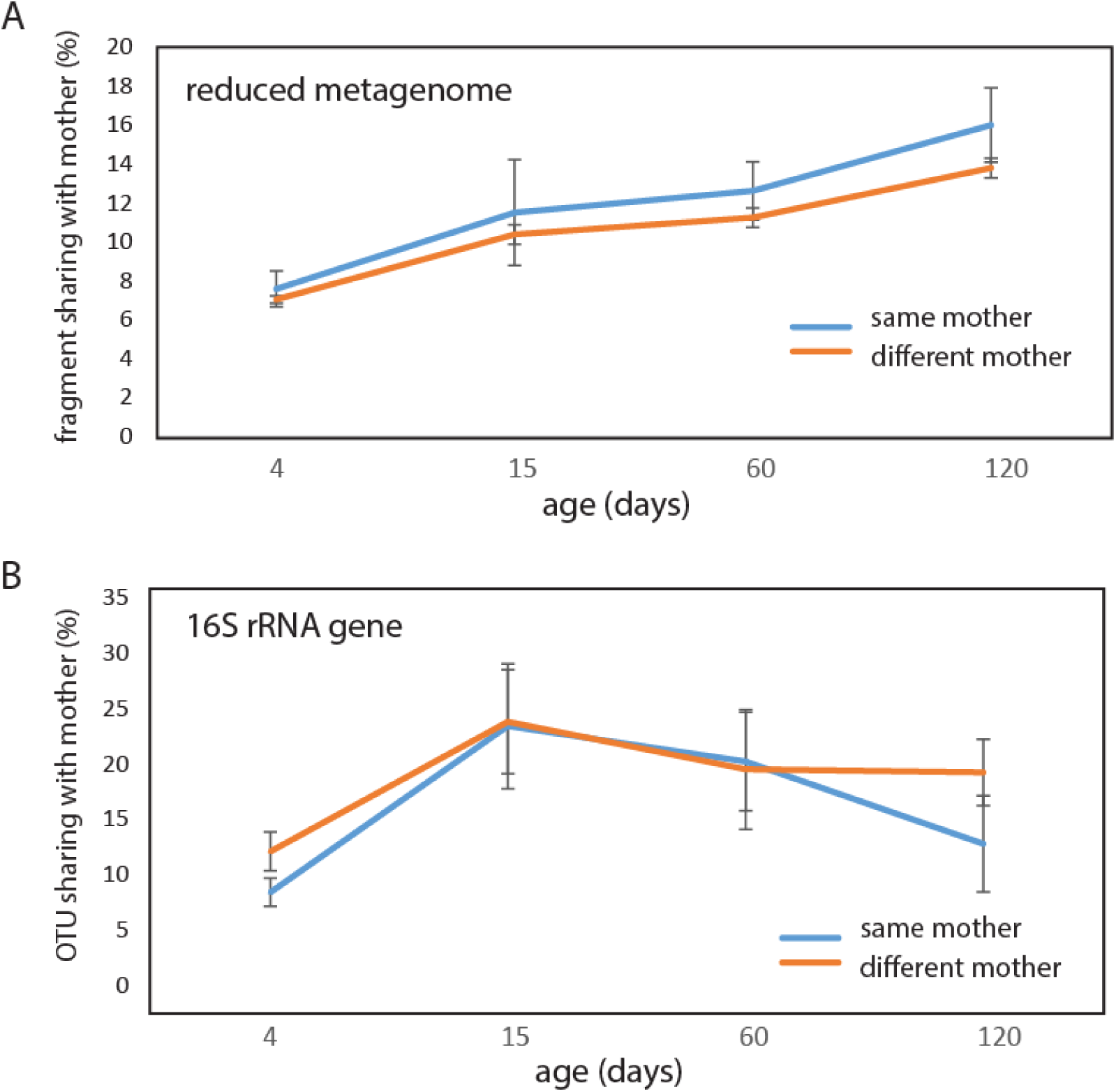
Sharing of microbiota with mothers for (A) RMS fragments and (B) 16S rRNA gene OTUs.

For the 16S rRNA gene sequence data we did not identify any significant association of OTUs with mode of delivery by ASCA-ANOVA. Furthermore, no significant association between 16S rRNA gene sequence data and antibiotic usage was determined.

### Vertical transmission

About 7% of the fragments detected by RMS for the 4-day-old children were shared with the mothers, with no difference between vaginally or c-section delivered children (p=0.67, Kruskal-Wallis test). Furthermore, there were no significant differences if the sharing was with the same or different mother for any of the age categories, although there was a tendency towards increased association with the same mother with age (Fig. 4A).

Similarly, as for RMS we did not identify any significant differences between the same or different mothers with respect to OTU sharing. However, the 16S rRNA gene OTUs displayed a pattern different from the reduced metagenome sequencing fragment sharing, with a peak in sharing at 15 days (Fig. 4B).

### Sharing within age categories

For the reduced metagenome sequencing, we found that the average sharing of fragments between individuals increased from below 15% for 4-day-old children to more than 30% for the mothers, with the 15-day to 4-month samples showing intermediate levels (Fig. 5A).

**Figure 5.**
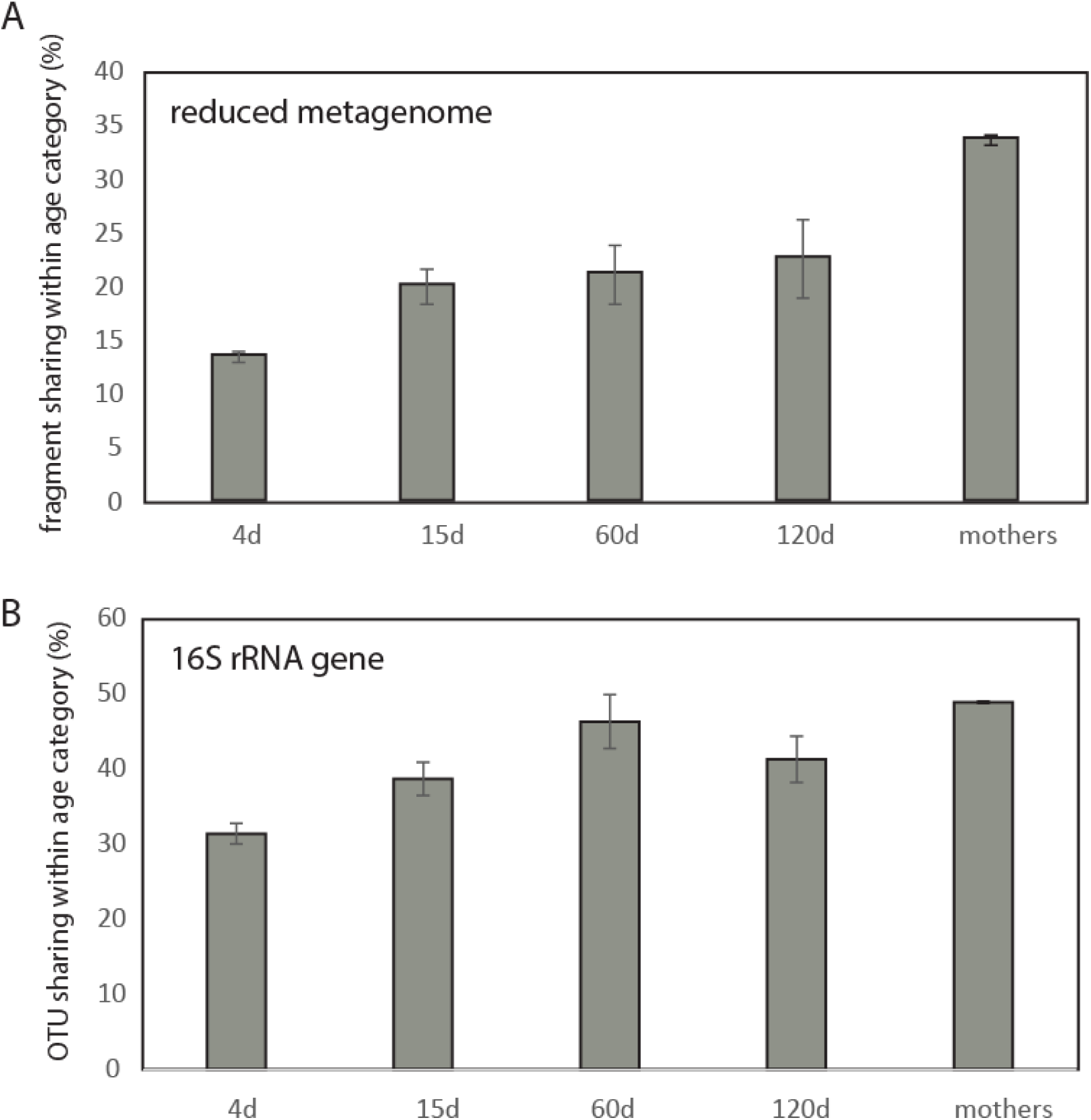
Sharing of microbiota with age categories for (A) RMS fragments and (B) 16S rRNA gene OTUs. Abbreviations d; days of life.

The age-related differences were less pronounced for the 16S rRNA gene sequence data, with an increase from 30% at 4 days to about 50% for the mothers. The 15-day to 4-month samples showed relatively large fluctuations for the shared 16S rRNA gene OTUs (Fig. 5B).

### Identification of core genomes

By RMS we identified the mothers’ core genomes, and the relative abundance of these genomes in infants. We first identified the RMS fragments shared across more than half of the mothers. In total, 10 009 RMS fragments satisfied this criterion (Fig. 2). From these, we identified 15 genome-sequenced species with more than 97% identity to the core fragments by Blast search (Fig. 2). The prevalence of these genomes in both mothers and 4-day-old children were determined by mapping all the RMS fragments, using the core genomes as reference. In mothers we identified 5 core genomes with a relative abundance above 1%, while for children only *Bacteroides vulgatus* showed high relative abundance (Fig. 6).

**Figure 6.**
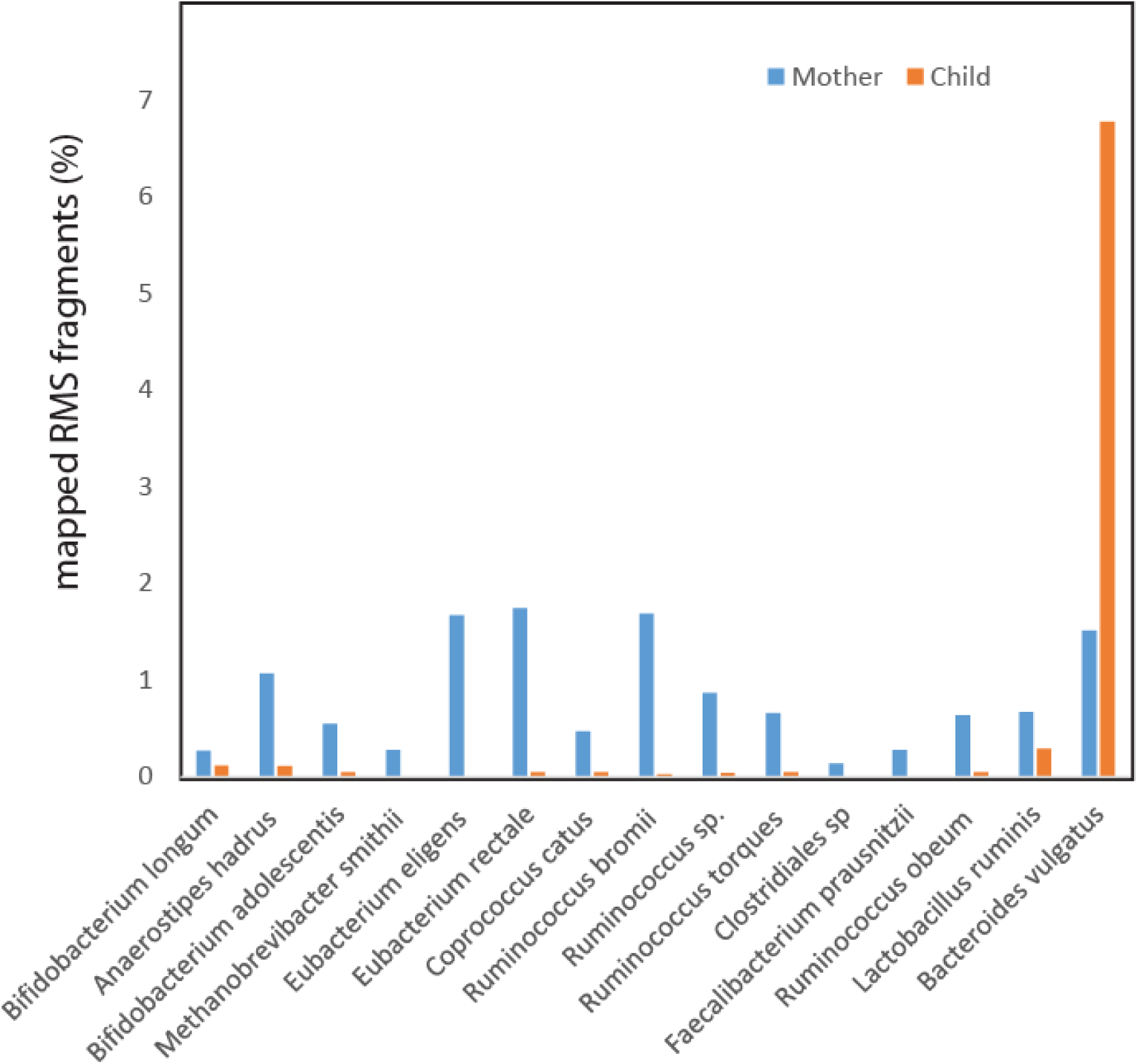
Sharing of core RMS fragments with genome sequenced prokaryotes.

### Validation of RMS

We validated RMS on experimental communities with known composition. This validation showed that there were clear signatures separating the bacteria in mixed populations, even between closely related *Bifidobacteria* (Suppl. Fig. 2A). Next we determined the quantitative potential of the RMS approach. This was done through regression analyses between the expected and observed DNA quantity of the four bacteria in the experimental community. All four evaluated bacteria showed high correlations (R2 > 0.8) between estimated concentrations based on RMS fragment frequency, and the expected concentrations (Suppl. Fig. 2B).

Finally, we determined the frequency of the reduced metagenome sequencing fragments from assembled shotgun data. This comparison showed a high correlation between the contig size and number of RMS fragments mapping to the respective contigs (Suppl. Fig. 3), with the mean distance between the RMS fragments being 5513±187 bp.

## DISCUSSION

There was a consistent increase in bacterial species shared between mothers and children with age based on the RMS data, while 16S rRNA gene analyses suggested a less consistent age-related pattern. This could be due to the fact that 16S rRNA gene analyses may merge several strains into the same OTUs (11), obscuring the analyses. For the mothers, the shotgun sequencing was far too shallow to yield any reasonable estimate of metagenome sequence size or strain composition, illustrating the shotgun sequencing challenges. The current approaches to extract strain level information from shotgun would require very deep sequencing (12). Therefore, we believe that the RMS approach can be a valuable and cost efficient contribution in deducing patterns associated with the gut microbiota.

Our RMS results support direct mother to child transmission of less than 10 % of stool-associated bacteria. Although the sharing was slightly higher with the child’s own mother rather than a random mother, most of the bacteria seem to be recruited from a common pool of gut associated bacteria. This contrasts with recent findings that suggest high strain sharing between infants and their mothers (4). However, given that more than one-third of the strains are shared across mothers, it would be difficult to determine if a strain is transmitted to a child from his or her mother or from some other adult. From the taxonomic identity, however, fragments belonging to the genus *Bacteroides* seemed underrepresented for children delivered by cesarean section. This is consistent with previous observations with long-term underrepresentation of *Bacteroides* in c-section delivered babies (13). The very high relative abundance of *B. vulgatus* in children could indicate that this bacterium plays an important role in the early development of the gut microbiota. *B. vulgatus* is mucin degrading bacterium (14) interacting with *Escherichia coli* in inflammation induction (15), and is suppressed by *Bifidobacteria* (16). To our knowledge, however, no studies have yet addressed the role of *B. vulgatus* in infants.

In our study we found very low levels of clostridia for the 4-day-old children, in addition to a lack of direct mother-child associations. Therefore, we found it unlikely that most of the adult associated clostridia are transmitted at delivery. Recently, it has been found that a large portion of gut bacteria are spore-formers (17), with endospores as a potential vector for transmission at a later stage than delivery (18).

Previous 16S rRNA gene sequencing have shown high degree of sharing at the genus/species level across mothers (3,19). Thus, the increased resolution of the reduced metagenome sequencing further supports the sharing of a relatively limited number of bacterial species/lineages within human populations. Our results suggest that one-third of the fragments are shared across random mothers. The mapping of the core fragments identified among the Indian mothers to human-derived genome sequenced isolates (mostly from Europe and America) further support the fact that there are limited number of human gut associated bacteria, and that these have wide geographic distribution. Interestingly, *Ruminococcus bromii,* which was among the most prevalent and dominant species for the Indian mothers, has previously been identified as a keystone species in resistant starch degradation, supporting the growth of both *Eubacterium rectale* and *Bifidobacterium adolescentis* (20), which were all identified among the 15 bacterial species shared across more than half of the Indian mothers in our work. This suggests that the core bacteria could have biologically important interactions.

Antibiotic usage during labor seemed to have a major impact on the resistance genes in the children without impacting the overall microbiota composition. This is consistent with previous observations suggesting that the mobilome can evolve independently of the overall composition of the gut microbiota (3). Furthermore, we detected resistance associations for antibiotics other than those used, indicating potential antibiotic resistance linkage (21). Thus, antibiotic usage during labor could be a major contributing factor to antibiotic resistance spread within the infants’ commensal gut microbiota.

## CONCLUSION

In conclusion, our results support a model with late recruitment of adult gut associated bacteria in infants, with a more than five-fold increase in genetic richness from child to adult.

## DECLARATIONS

### Availability of data and material

The raw sequencing reads are deposited in the European Nucleotide Archive with the accession number PRJEB85416, while the data used for figure generation are provided in a Supplementary Excel file.

### Ethics approval and consent to participate

A written consent was obtained from all the participants

### Consent for publication

Not applicable.

### Author’s contributions

AR designed the study. EA did the methods validation. IA performed the analyses. JL did the shotgun analyses. PM, SP and RN did the sample collection and recording of metadata. KR analyzed the data, wrote the paper and invented the RMS methods. All authors commented on the manuscript.

## Acknowledgements

We would like to thank the Norwegian government for the scholarship provided to Anuradha Ravi. We would also like to thank the doctors and nurses at Nishanth Hospital, India for doing the sampling and collecting the information.

## SUPPLEMENTARY INFORMATION

**Supplementary Table 1.**
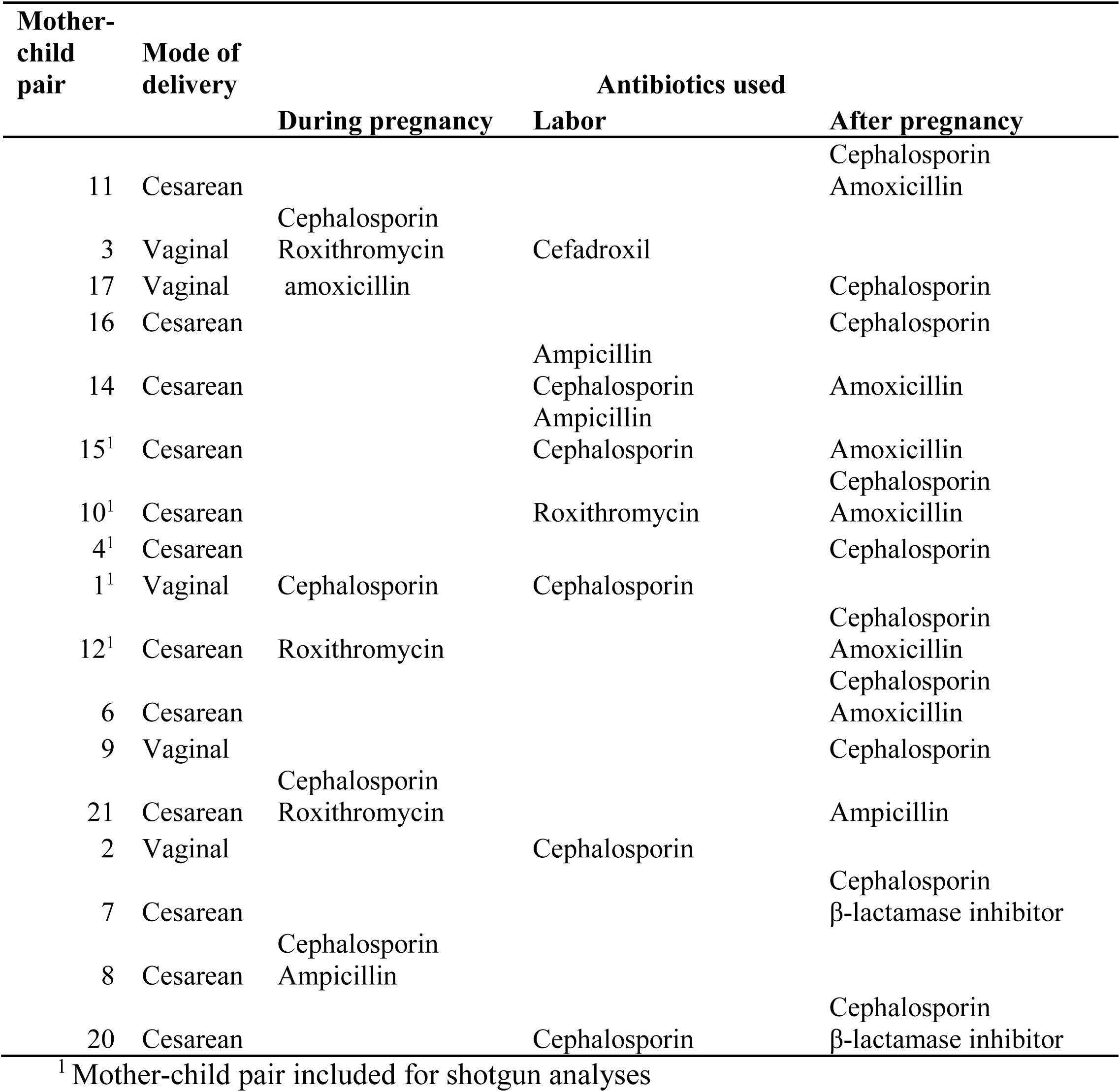
Metadata of the mother-child pairs.

**Supplementary Table 2.**
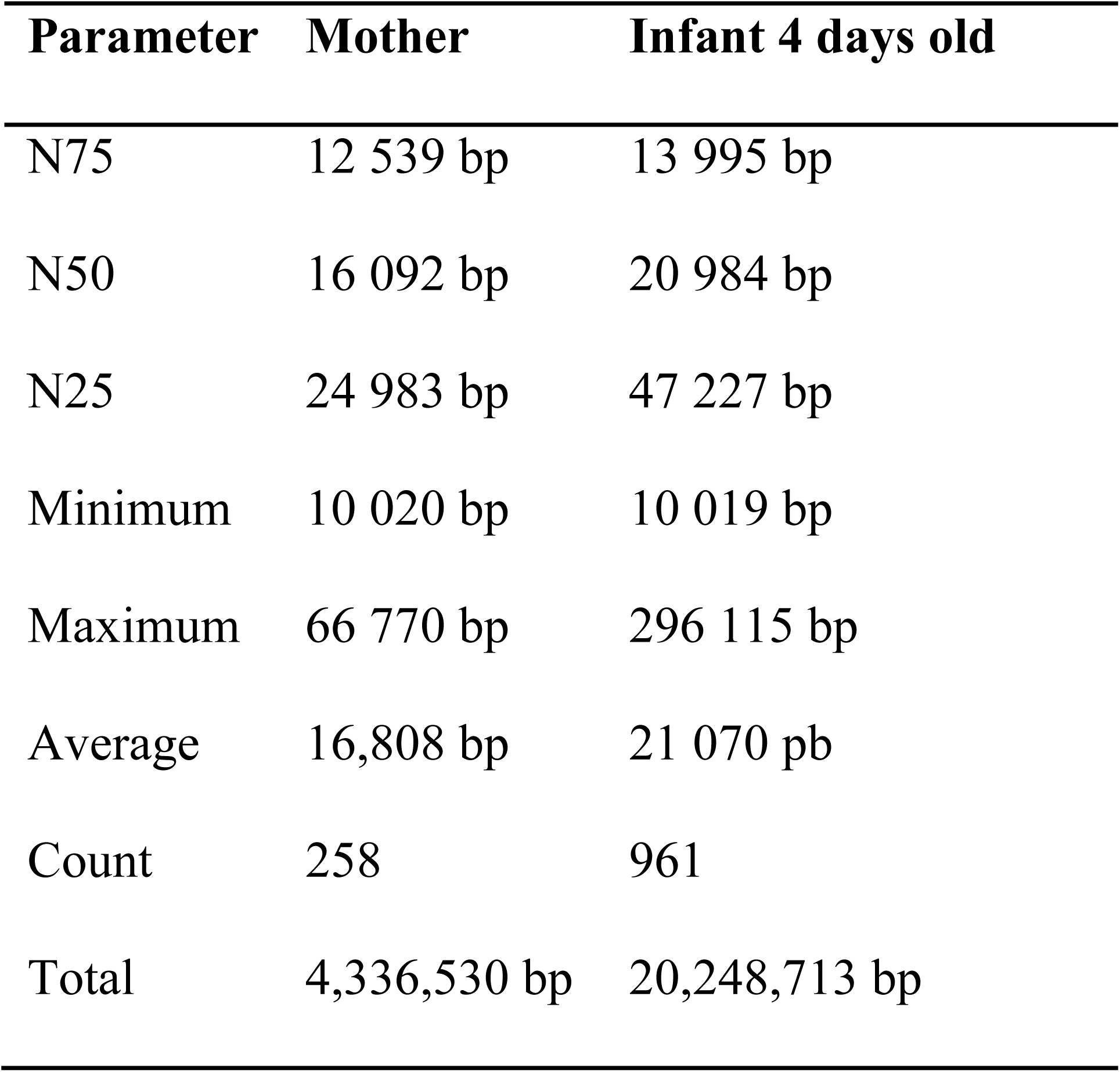
Metagenome assembly parameters.

**Supplementary Table 3.**
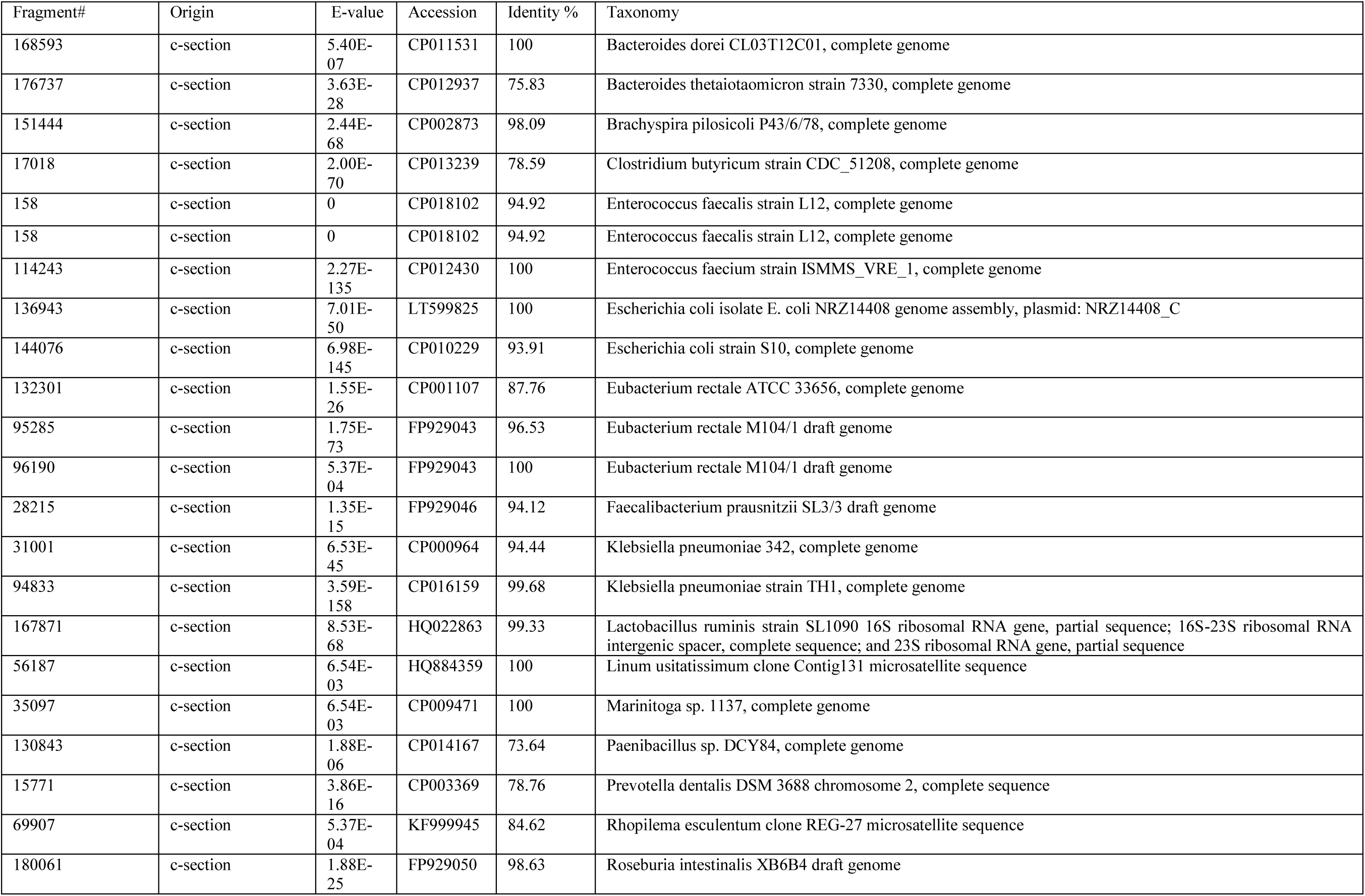
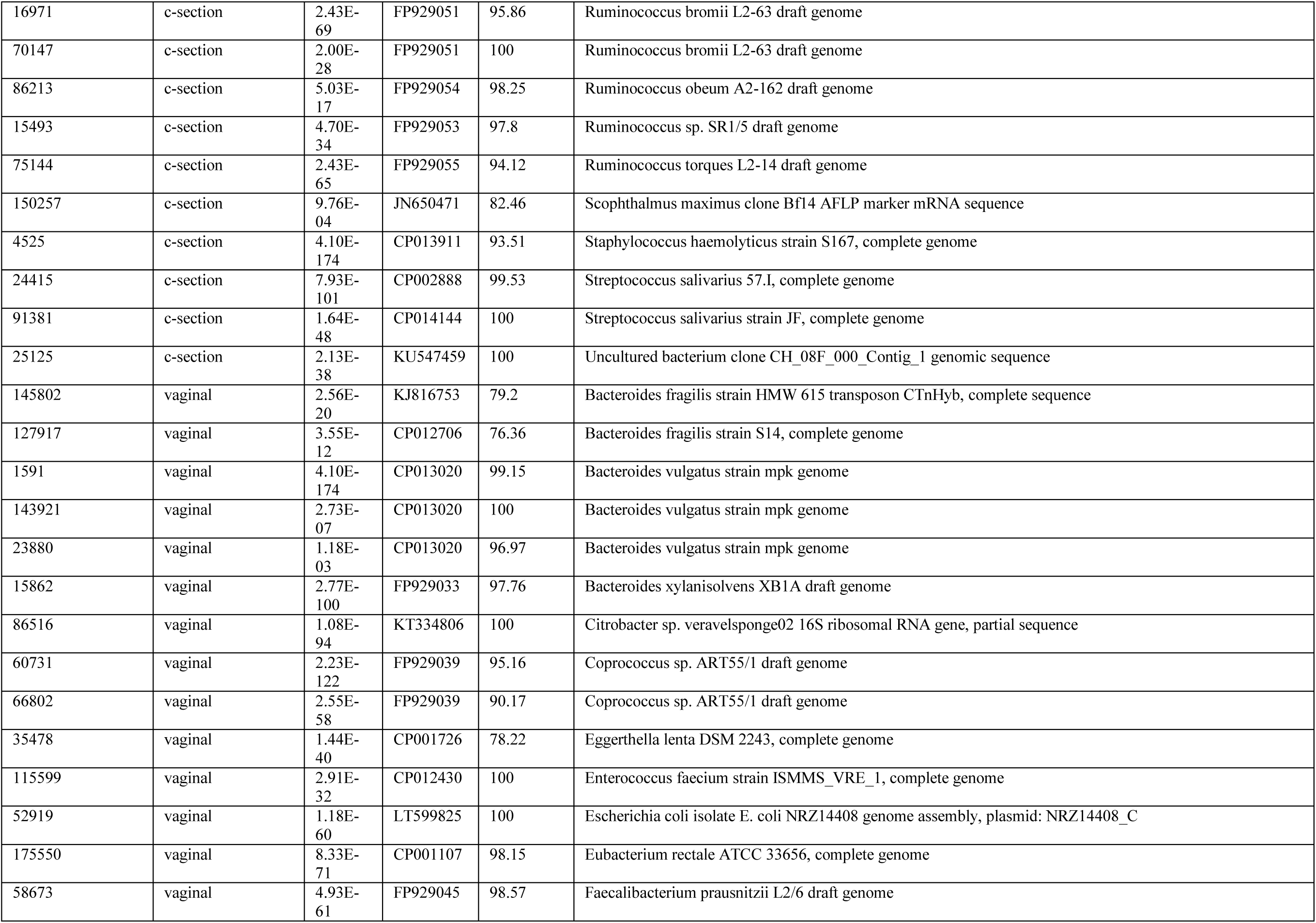
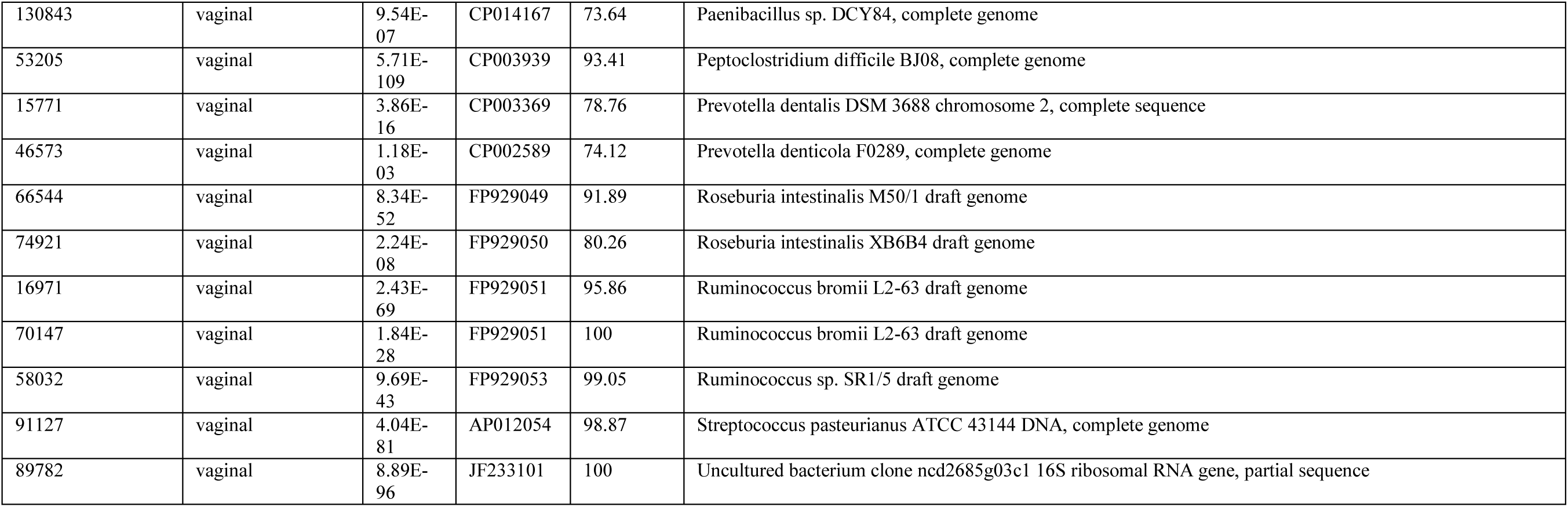
Fragments associated with vaginal delivery and c-section.

| Fragment# | Origin | E-value | Accession | Identity % | Taxonomy |
| --- | --- | --- | --- | --- | --- |
| 168593 | c-section | 5.40E-07 | CP011531 | 100 | Bacteroides dorei CL03T12C01, complete genome |
| 176737 | c-section | 3.63E-28 | CP012937 | 75.83 | Bacteroides thetaiotaomicron strain 7330, complete genome |
| 151444 | c-section | 2.44E-68 | CP002873 | 98.09 | Brachyspira pilosicoli P43/6/78, complete genome |
| 17018 | c-section | 2.00E-70 | CP013239 | 78.59 | Clostridium butyricum strain CDC_51208, complete genome |
| 158 | c-section | 0 | CP018102 | 94.92 | Enterococcus faecalis strain L12, complete genome |
| 158 | c-section | 0 | CP018102 | 94.92 | Enterococcus faecalis strain L12, complete genome |
| 114243 | c-section | 2.27E-135 | CP012430 | 100 | Enterococcus faecium strain ISMMS_VRE_1, complete genome |
| 136943 | c-section | 7.01E-50 | LT599825 | 100 | Escherichia coli isolate E. coli NRZ14408 genome assembly, plasmid: NRZ14408_C |
| 144076 | c-section | 6.98E-145 | CP010229 | 93.91 | Escherichia coli strain S10, complete genome |
| 132301 | c-section | 1.55E-26 | CP001107 | 87.76 | Eubacterium rectale ATCC 33656, complete genome |
| 95285 | c-section | 1.75E-73 | FP929043 | 96.53 | Eubacterium rectale M104/1 draft genome |
| 96190 | c-section | 5.37E-04 | FP929043 | 100 | Eubacterium rectale M104/1 draft genome |
| 28215 | c-section | 1.35E-15 | FP929046 | 94.12 | Faecalibacterium prausnitzii SL3/3 draft genome |
| 31001 | c-section | 6.53E-45 | CP000964 | 94.44 | Klebsiella pneumoniae 342, complete genome |
| 94833 | c-section | 3.59E-158 | CP016159 | 99.68 | Klebsiella pneumoniae strain TH1, complete genome |
| 167871 | c-section | 8.53E-68 | HQ022863 | 99.33 | Lactobacillus ruminis strain SL1090 16S ribosomal RNA gene, partial sequence; 16S-23S ribosomal RNA intergenic spacer, complete sequence; and 23S ribosomal RNA gene, partial sequence |
| 56187 | c-section | 6.54E-03 | HQ884359 | 100 | Linum usitatissimum clone Contig131 microsatellite sequence |
| 35097 | c-section | 6.54E-03 | CP009471 | 100 | Marinitoga sp. 1137, complete genome |
| 130843 | c-section | 1.88E-06 | CP014167 | 73.64 | Paenibacillus sp. DCY84, complete genome |
| 15771 | c-section | 3.86E-16 | CP003369 | 78.76 | Prevotella dentalis DSM 3688 chromosome 2, complete sequence |
| 69907 | c-section | 5.37E-04 | KF999945 | 84.62 | Rhopilema esculentum clone REG-27 microsatellite sequence |
| 180061 | c-section | 1.88E-25 | FP929050 | 98.63 | Roseburia intestinalis XB6B4 draft genome |
| 16971 | c-section | 2.43E-69 | FP929051 | 95.86 | Ruminococcus bromii L2-63 draft genome |
| 70147 | c-section | 2.00E-28 | FP929051 | 100 | Ruminococcus bromii L2-63 draft genome |
| 86213 | c-section | 5.03E-17 | FP929054 | 98.25 | Ruminococcus obeum A2-162 draft genome |
| 15493 | c-section | 4.70E-34 | FP929053 | 97.8 | Ruminococcus sp. SR1/5 draft genome |
| 75144 | c-section | 2.43E-65 | FP929055 | 94.12 | Ruminococcus torques L2-14 draft genome |
| 150257 | c-section | 9.76E-04 | JN650471 | 82.46 | Scophthalmus maximus clone Bf14 AFLP marker mRNA sequence |
| 4525 | c-section | 4.10E-174 | CP013911 | 93.51 | Staphylococcus haemolyticus strain S167, complete genome |
| 24415 | c-section | 7.93E-101 | CP002888 | 99.53 | Streptococcus salivarius 57.I, complete genome |
| 91381 | c-section | 1.64E-48 | CP014144 | 100 | Streptococcus salivarius strain JF, complete genome |
| 25125 | c-section | 2.13E-38 | KU547459 | 100 | Uncultured bacterium clone CH_08F_000_Contig_1 genomic sequence |
| 145802 | vaginal | 2.56E-20 | KJ816753 | 79.2 | Bacteroides fragilis strain HMW 615 transposon CTnHyb, complete sequence |
| 127917 | vaginal | 3.55E-12 | CP012706 | 76.36 | Bacteroides fragilis strain S14, complete genome |
| 1591 | vaginal | 4.10E-174 | CP013020 | 99.15 | Bacteroides vulgatus strain mpk genome |
| 143921 | vaginal | 2.73E-07 | CP013020 | 100 | Bacteroides vulgatus strain mpk genome |
| 23880 | vaginal | 1.18E-03 | CP013020 | 96.97 | Bacteroides vulgatus strain mpk genome |
| 15862 | vaginal | 2.77E-100 | FP929033 | 97.76 | Bacteroides xylanisolvens XB1A draft genome |
| 86516 | vaginal | 1.08E-94 | KT334806 | 100 | Citrobacter sp. veravelsponge02 16S ribosomal RNA gene, partial sequence |
| 60731 | vaginal | 2.23E-122 | FP929039 | 95.16 | Coprococcus sp. ART55/1 draft genome |
| 66802 | vaginal | 2.55E-58 | FP929039 | 90.17 | Coprococcus sp. ART55/1 draft genome |
| 35478 | vaginal | 1.44E-40 | CP001726 | 78.22 | Eggerthella lenta DSM 2243, complete genome |
| 115599 | vaginal | 2.91E-32 | CP012430 | 100 | Enterococcus faecium strain ISMMS_VRE_1, complete genome |
| 52919 | vaginal | 1.18E-60 | LT599825 | 100 | Escherichia coli isolate E. coli NRZ14408 genome assembly, plasmid: NRZ14408_C |
| 175550 | vaginal | 8.33E-71 | CP001107 | 98.15 | Eubacterium rectale ATCC 33656, complete genome |
| 58673 | vaginal | 4.93E-61 | FP929045 | 98.57 | Faecalibacterium prausnitzii L2/6 draft genome |
| 130843 | vaginal | 9.54E-07 | CP014167 | 73.64 | Paenibacillus sp. DCY84, complete genome |
| 53205 | vaginal | 5.71E-109 | CP003939 | 93.41 | Peptoclostridium difficile BJ08, complete genome |
| 15771 | vaginal | 3.86E-16 | CP003369 | 78.76 | Prevotella dentalis DSM 3688 chromosome 2, complete sequence |
| 46573 | vaginal | 1.18E-03 | CP002589 | 74.12 | Prevotella denticola F0289, complete genome |
| 66544 | vaginal | 8.34E-52 | FP929049 | 91.89 | Roseburia intestinalis M50/1 draft genome |
| 74921 | vaginal | 2.24E-08 | FP929050 | 80.26 | Roseburia intestinalis XB6B4 draft genome |
| 16971 | vaginal | 2.43E-69 | FP929051 | 95.86 | Ruminococcus bromii L2-63 draft genome |
| 70147 | vaginal | 1.84E-28 | FP929051 | 100 | Ruminococcus bromii L2-63 draft genome |
| 58032 | vaginal | 9.69E-43 | FP929053 | 99.05 | Ruminococcus sp. SR1/5 draft genome |
| 91127 | vaginal | 4.04E-81 | AP012054 | 98.87 | Streptococcus pasteurianus ATCC 43144 DNA, complete genome |
| 89782 | vaginal | 8.89E-96 | JF233101 | 100 | Uncultured bacterium clone ncd2685g03c1 16S ribosomal RNA gene, partial sequence |

**Supplementary Table 4.**
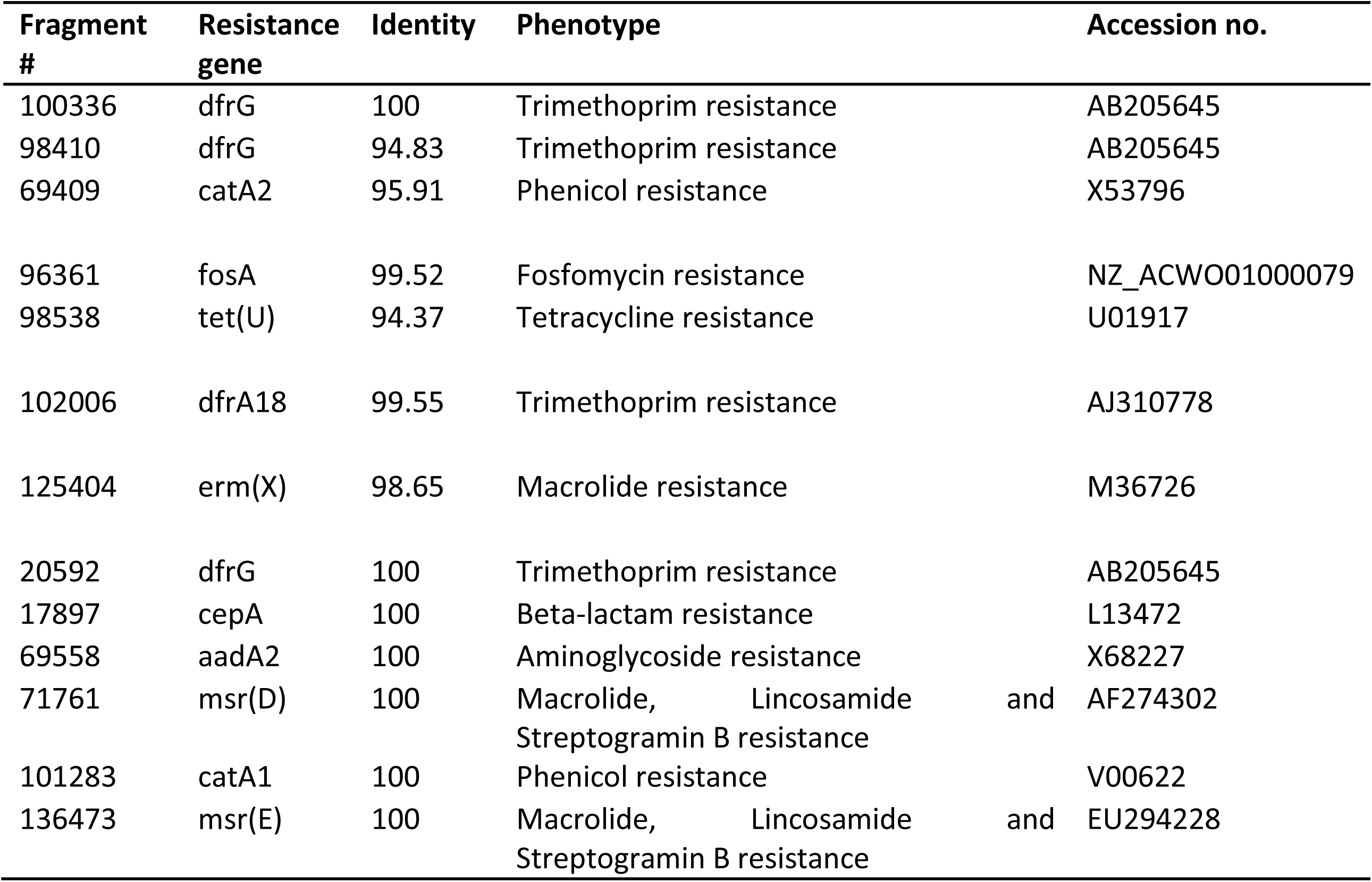
Antibiotic resistance genes detected by reduced metagenome sequencing.

**Suppl. Fig. 1.**
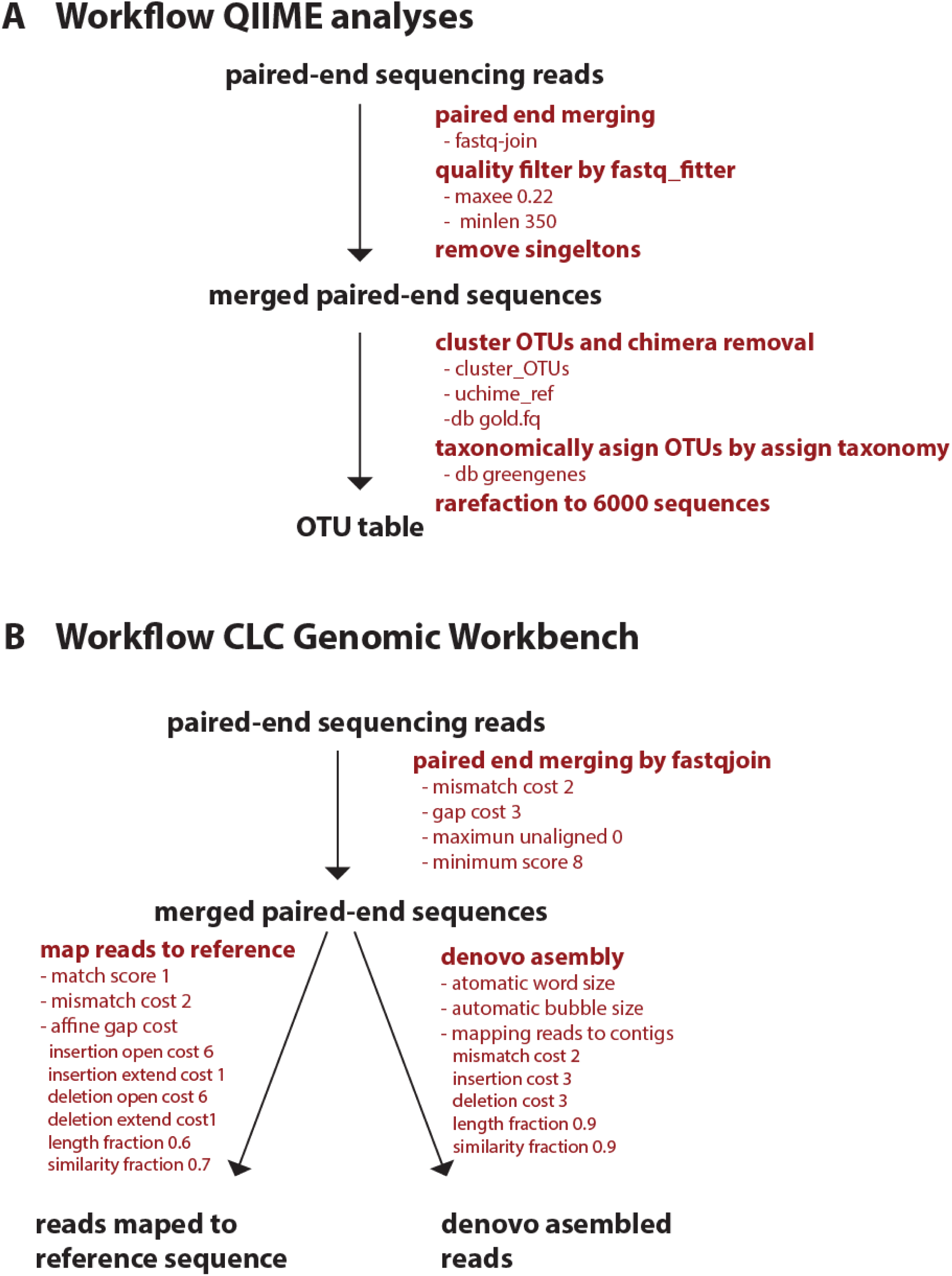
Workflow for QIIME analyses (A), and CLC Genomic Workbench (B).

**Suppl. Fig. 2.**
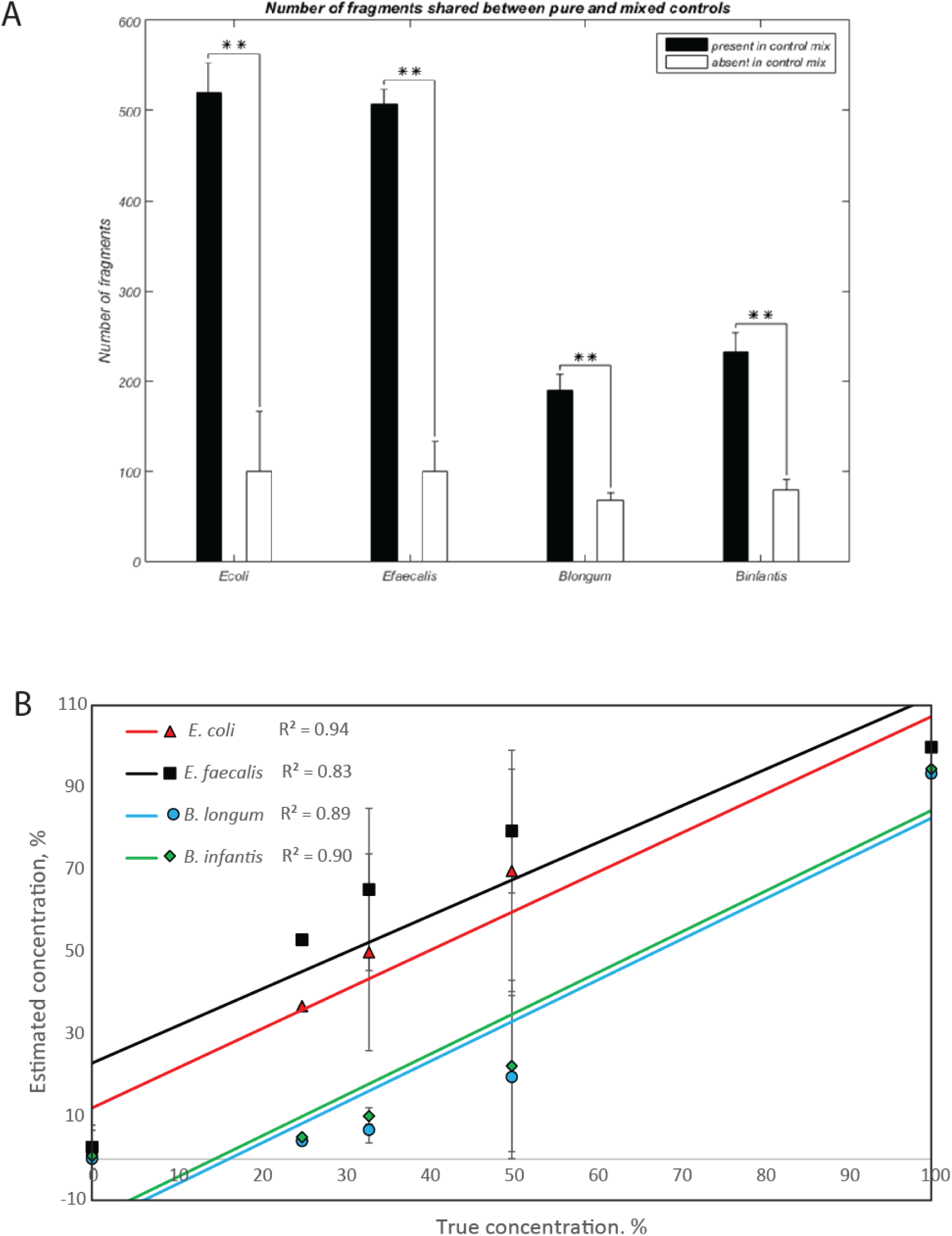
Evaluation of (A) the uniqueness of the reduced metagenome fragments and (B) the quantitative properties. The true concentrations are based amount of DNA added for the different species.

**Suppl. Fig. 3.**
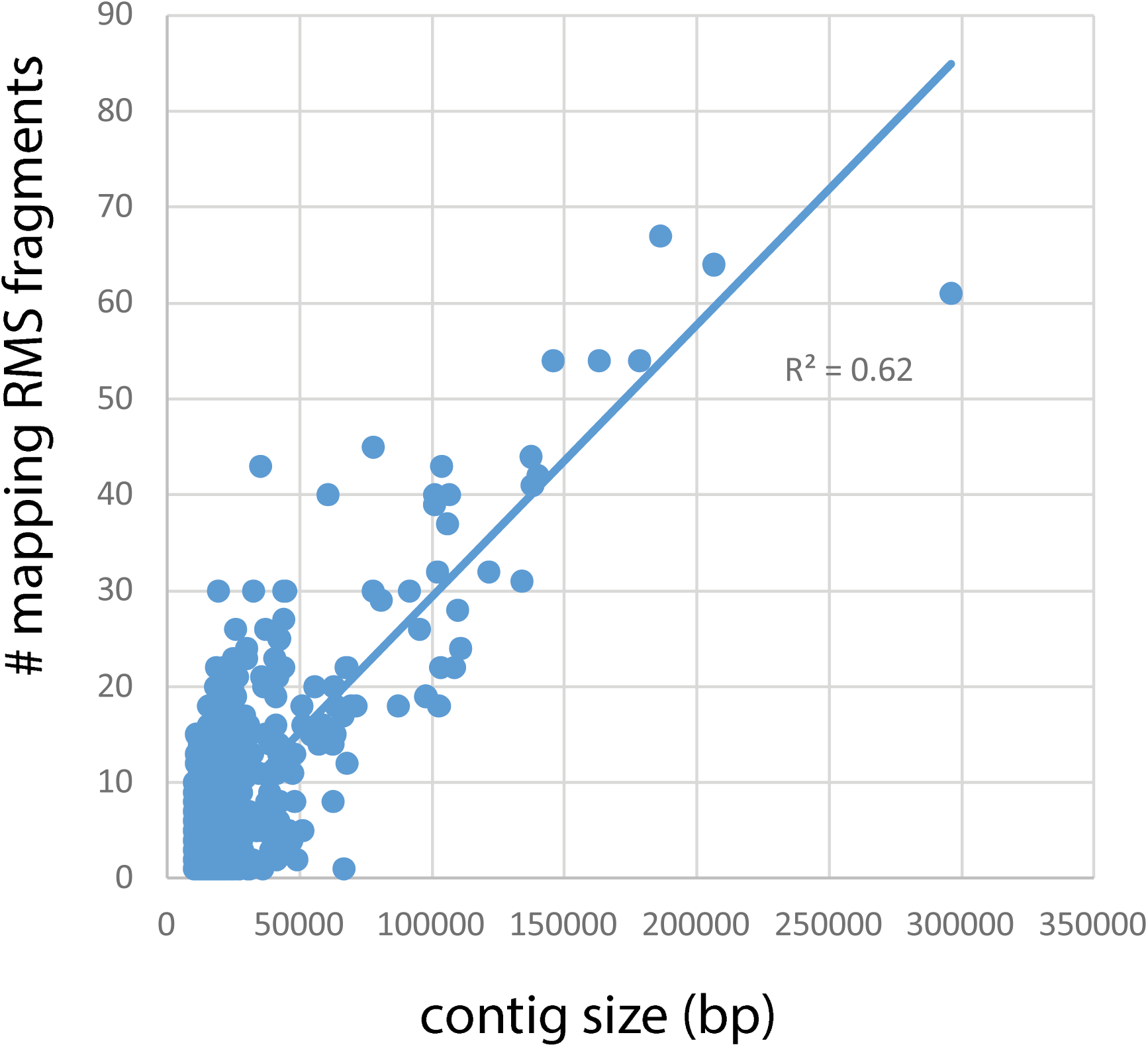
Correlation between number of RMS fragments detected and contig size. The number of mapping fragments was determined by using the contigs as reference for RMS fragment mapping.

**Suppl. Fig. 4.**
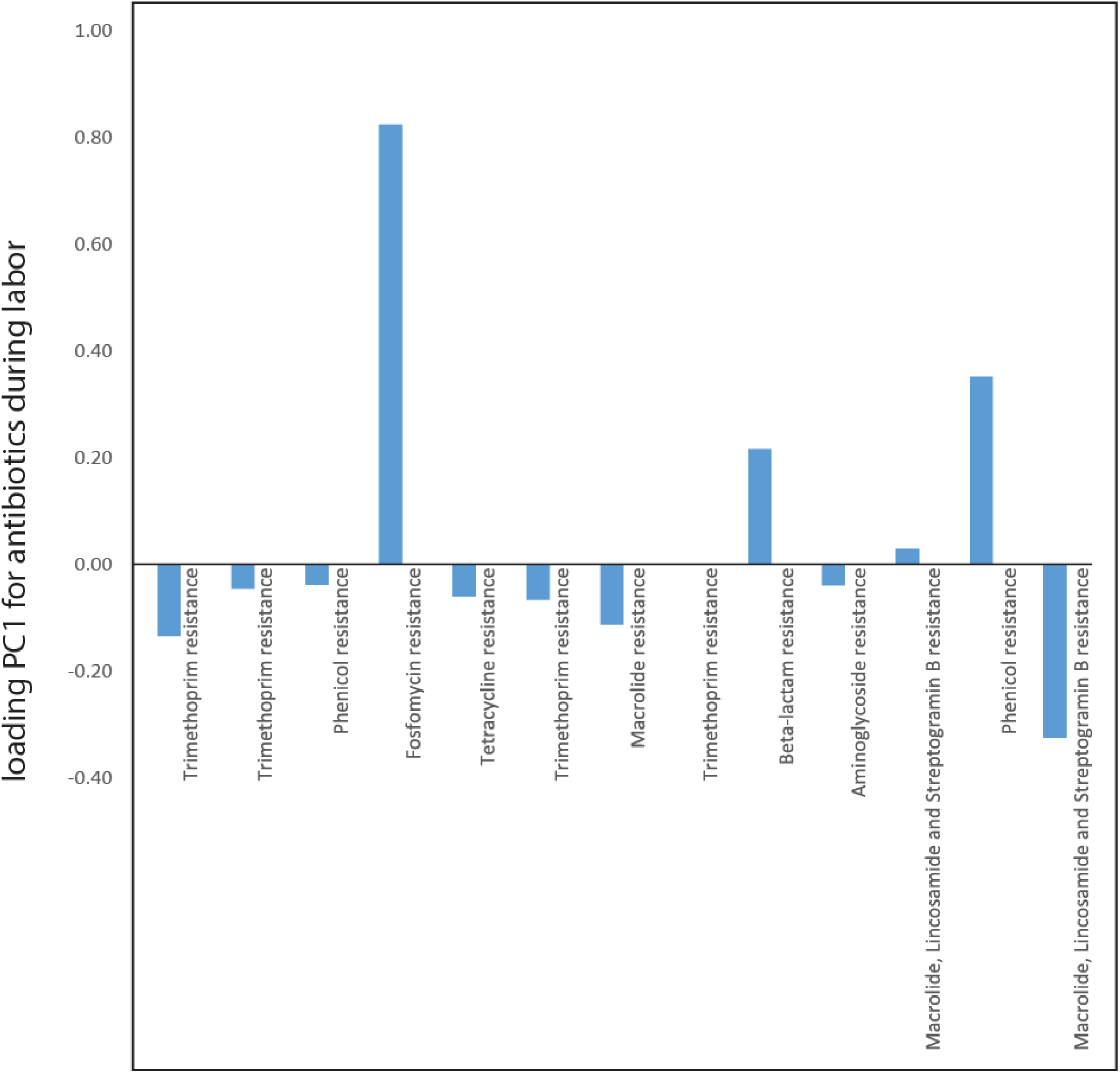
Antibiotic resistance associated with antibiotic usage during labor. The importance (principal component loading) of the different resistance genes in explaining the overall association with antibiotic usage.

